# Ranking of Major Classes of Antibiotics for Activity against Stationary Phase *Pseudomonas aeruginosa* and Identification of Clinafloxacin + Cefuroxime + Gentamicin Drug Combination that Eradicates Persistent *P. aeruginosa* Infection in a Murine Cystic Fibrosis Model

**DOI:** 10.1101/686105

**Authors:** Yuting Yuan, Rebecca Yee, Naina Gour, Xinzhong Dong, Jie Feng, Wanliang Shi, Ying Zhang

## Abstract

*Pseudomonas aeruginosa* can cause serious persistent infections such as ventilator-associated pneumonia, sepsis, biofilm-related infections as in cystic fibrosis (CF) patients. Although CF lung infections can be treated with antibiotics, full clearance is difficult due to *P. aeruginosa* persistence. While antibiotic activity against growing *P. aeruginosa* is well documented, their activity against the non-growing persisters enriched in stationary phase cultures has not been well studied. Here, we systematically evaluated and ranked the six major classes of antibiotics, cell wall and cell membrane inhibitors, protein synthesis inhibitors, DNA synthesis inhibitors, RNA synthesis inhibitors, sulfa drugs, and nitrofurantoin, for their activity against both growing and persister forms of *P. aeruginosa* using colony forming count (CFU) and SYBR Green I/Propidium Iodide (PI) viability assay. Among the six major classes of antibiotics, cell wall and cell membrane inhibitors (Cefuroxime and Colistin), DNA synthesis inhibitors (Clinafloxacin) and sulfa drugs (Sulfamethoxazole) had good activity against stationary phase cells. In contrast, protein synthesis inhibitors (Gentamicin), RNA synthesis inhibitor (Rifampicin) and Nitrofurantoin had relatively poor activity against the stationary phase *P. aeruginosa* but relatively high activity against log phase *P. aeruginosa.* Clinafloxacin is the only single drug that could completely kill all (10^9^ CFU) stationary phase cells in a 4 day drug exposure. The Cefuroxime + Gentamicin+ Clinafloxacin combination could kill all biofilm bacteria in 2 days whereas the clinically used drug combination Cefuroxime + Gentamicin + Colistin only partially killed the biofilm bacteria with 10^3^ CFU remaining. In a murine persistent CF lung infection model, only Cefuroxime + Gentamicin+ Clinafloxacin cleared all bacteria in the infected lungs, whereas Clinafloxacin alone, or Cefuroxime + Clinafloxacin, or the current recommended drug combination Cefuroxime + Gentamicin, all failed to completely clear the bacterial load in the lungs. The complete sterilization of the bacterial load is a property of Clinafloxacin combination, as Cefuroxime + Gentamicin+ Levofloxacin combination was unable to clear the bacterial load in the lungs. Our findings demonstrate the importance of persister drug clinafloxacin, offer new therapeutic approaches for more effective treatment of persistent *P. aeruginosa* infections, and may have implications for treating other persistent infections.

## Introduction

*Pseudomonas aeruginosa* is a highly resistant, opportunistic Gram-negative bacterium (Stover et al. 2000) that causes serious infections in hospitalized patients or people with compromised immune systems. Patients with burn wounds, cystic fibrosis, acute leukemia, organ transplants, and intravenous-drug users are at high risk for infections (Aloush et al. 2006). Because *P. aeruginosa* causes many nosocomial infections, epidemics within the hospitals have been recorded (Bodey et al. 1983; Cross et al. 1983). *P. aeruginosa* is a major pathogen in the cystic fibrosis (CF) lung, and it causes a persistent infection that cannot be eradicated by even the most aggressive antibiotic therapy (Costerton, Stewart, and Greenberg 1999b; Hoiby 1993). This has been attributed to bacterial biofilms which are resistant or tolerant to antibiotic treatments and can evade host immune defense (Costerton, Stewart, and Greenberg 1999b; J W Costerton et al. 1995). Based on the Johns Hopkins Antibiotics Guide(Johns Hopkins Hospital 2017), high dose of synergistic antibiotic combinations (β-lactam + aminoglycoside) may improve outcomes of serious *P. aeruginosa* infections in immunocompromised hosts clinically and these combinations were determined against *Pseudomonas aeruginosa* from the sputum of patients with cystic fibrosis (Scribner et al. 1982). For multidrug resistant strains, colistin can be added to the above treatment (Florescu et al. 2012). Due to increase antibiotic resistance, the repertoire of effective agents against *P. aeruginosa* is limited (Rossolini and Mantengoli 2005), and thus better therapies are needed.

Bacterial cells may escape the effects of antibiotics due to epigenetic changes; these cells are known as persisters (Wood, Knabel, and Kwan 2013). Many chronic infections are associated with the ability of the bacteria to predominantly colonize body surfaces and tissues as multicellular aggregation as biofilms (Costerton, Stewart, and Greenberg 1999a; Wolfmeier et al. 2018). Formation of these sessile communities and their inherent resistance to antimicrobial agents are at the root of many persistent and chronic bacterial infections. The number of persisters in a growing population of bacteria rises at mid-log and reaches a maximum of approximately 1% at stationary state (Lewis 2008). Similarly, slow-growing biofilms produce substantial numbers of persisters. The ability of a biofilm to limit the access of the immune system components, and the ability of persisters to sustain an antibiotic attack could then account for the recalcitrance of such infections *in vivo* and frequent relapse after treatment.

The most common method to assess the killing activity of antibiotics against stationary phase bacteria is through counting of viable cells grown on agar plates in colony forming unit (CFU) assay. A major disadvantage of CFU counting is the lengthy time (1-3 days) for bacteria to grow on agar plates and miss the subpopulation of viable but non-culturable bacteria that do not form CFUs. To more rapidly quantify the amount of live cells after drug treatment, a SYBR Green I/Propidium Iodide (PI) viability assay (Klockgether et al. 2010) was developed to evaluate antibiotic susceptibility and to perform high-throughput drug screens in *B. burgdorferi* (Feng et al. 2014). SYBR Green I is a high affinity dye that binds double-stranded DNA (dsDNA) and is commonly used to stain nucleic acids in PCR and flow cytometric analysis (Simpson et al. 2000; Morrison, Weis, and Wittwer 1998; Barbesti et al. 2000). SYBR Green I is a permeable dye that stains all live cells green, whereas PI is an impermeable dye that stains dead or damaged cells with compromised cell membrane red (Nicoletti et al. 1991). Therefore, fluorescence microscopy or fluorescence microplate readers can be used to measure the live/dead ratio of a bacterial sample as a rapid method to assess bacterial viability after drug treatment without CFU counts. Here, we applied the SYBR Green I/PI viability assay to *P. aeruginosa* and ranked six major classes of antibiotics in killing growing and stationary non-growing forms of *P. aeruginosa* compared with conventional CFU based viability assay. Using these methods, we also identified drug combinations that could more effectively eradicate persisters in stationary phase culture of *P. aeruginosa* as well as in biofilms *in vitro* and more importantly, in a persistent lung infection mouse model *in vivo*.

## Materials and Methods

### Bacterial strain, culture media and growth condition

*P. aeruginosa* strain PAO-1 was cultured in tryptic soy broth (TSB) medium. The culture was incubated at 37℃ with shaking at 200 rpm. After overnight incubation, stationary phase cultures were exposed to different antibiotics and incubated at indicated times before washing and plating on tryptic soy agar (TSA) plates (Becton Dickinson) for colony forming unit (CFU) enumeration. The plates were incubated at 37 ℃ overnight.

### Preparation of antibiotics

Stock solutions of antibiotics Cefuroxime, Colistin, Gentamicin, Clinafloxacin, Rifampicin, Sulfamethoxazole and Nitrofurantoin (Sigma Aldrich Co.) were prepared with appropriate solvents at 10 mg/ml. Each antibiotic (25 μM) was added into 500 μL of stationary phase culture of *P. aeruginosa* and was incubated at 37 ℃ without shaking for various times. In addition, appropriate concentrations of the antibiotics were used for MIC determination using the microdilution method in 96-well plates (see below).

### Drug exposure assay by CFU counting on plates

Viable bacterial cells were determined by CFU count after drug exposure at days 2, 4, 6 and 10 as described (Harley 2008). First, bacterial suspensions (50 μL) after drug exposure were washed with fresh TSB medium twice and serial dilutions were prepared. 10 μL of each dilution was dropped onto TSA plates in triplicate followed by incubation at 37 ℃ overnight. The CFU/ml was calculated accordingly.

### SYBR Green I/PI assay for *P. aeruginosa*

The staining dyes were prepared by mixing SYBR Green I/PI (10,000X stock, Invitrogen) with propidium iodide (20 mM, Sigma-Aldrich) in distilled water and the ratio of SYBR Green I to propidium iodide was 1:3 in 100 µl distilled H_2_O(Feng et al. 2014). The drug treated samples were aliquoted and diluted 1:50 with fresh TSB medium. The SYBR Green I/PI dye mix (10 µL) was added to each 100 µL of sample. Each sample was vortexed and incubated at room temperature in the dark for 20 minutes. After incubation, each sample was transferred into 96-well plate. With excitation wavelength of 485 nm and 538 nm and 612 nm for green and red emission, respectively, the green and red fluorescence intensity was determined for each sample using a Synergy H1 microplate reader by BioTek Instruments (VT, USA). To generate the regression equation and regression curve of the relationship between the percentage of live bacteria and green/red fluorescent ratios, different proportions of live and 70% isopropyl alcohol killed dead cells were prepared (0:10, 1:4, 5:5, 4:1, 10:0) and both live and dead samples were diluted 50 times with fresh TSB medium first. Each proportion of live/dead *P. aeruginosa* was mixed with SYBR Green I/PI dye into each well of 96-well plate and the green/red fluorescence ratios were measured as described above, generating standard curve and equation with least-square fitting analysis. We used this equation to calculate the percentage of live cells of *P. aeruginosa*. Fluorescence microscopy imaging visualizing live and dead cells was performed using a Keyence BZ-X710 Fluorescence Microscope and was processed by BZ-X Analyzer provided by Keyence (Osaka, Japan).

### The MIC (minimum inhibitory concentration) determination

The standard microdilution method was used to determine the MIC of each antibiotic as described (Andrews 2001; Lambert 2002; Wiegand, Hilpert, and Hancock 2008). *P. aeruginosa* (1×10^9^) was inoculated (10 μL) into each well of 96-well plate containing 90 μL fresh TSB medium per well. Then each antibiotic was added into the well and the serial dilutions of drug treatment were made from 16, 8, 4, 2, 1 and 0 μg/mL. All experiments were run in duplicate or triplicate. The 96-well plate was incubated in 37 ℃ incubator overnight. The MIC is the lowest concentration of the antibiotic that prevented visible growth of *P. aeruginosa*.

### *P. aeruginosa* biofilm preparation

The *P. aeruginosa* biofilm model was prepared based on the protocol as described (Merritt, Kadouri, and O’Toole 2005). We first inoculated *P. aeruginosa* PAO-1 bacteria strain in a 5 ml fresh TSB culture and let it grow to stationary phase overnight. Then this stationary phase culture was diluted 1:100 in fresh TSB medium, and 100 μl diluted culture was pipetted into each well of 96-well plate and then the covered plate was put into 37 ℃ incubator overnight and *P. aeruginosa* biofilm attached on the bottom of the plate after removing the supernatant medium by pipetting on the side of the well while leaving the biofilm formed at the bottom of the plate intact.

### Drug combination assay on *P. aeruginosa* stationary phase persister model and biofilm model *in vitro*

Based on the Johns Hopkins Hospital Antibiotics Guideline (Johns Hopkins Hospital 2017), using high doses of synergistic antibiotic combinations (β-lactam + aminoglycoside) is considered to be able to improve outcomes of serious infections in immunocompromised hosts (Martinez et al. 2010). As for the multidrug resistant strains, colistin can also be added to the above double combination (Florescu et al. 2012). We evaluated Clinafloxacin (1.5 μg/ml) in combination with Cefuroxime (Cmax: 5 μg/ml), Gentamicin (Cmax: 10 μg/ml) and Colistin (Cmax: 5 μg/ml) separately. Colistin was replaced by Clinafloxacin in the clinically used triple drug combination Cefuroxime + Gentamicin + Colistin, we. The designed drug combinations or their single or two drug controls were added directly to stationary phase culture and CFU count was performed after 2 day and 4 day drug treatment. For the biofilm model, the designed drug combinations were prepared with MOPS buffer (1X) (diluted from 10X MOPS from Sigma Aldrich) to the final drug concentration and then transferred in to 96-well plate with *P. aeruginosa* biofilm attached to the bottom. Biofilm was washed with phosphate buffered saline (PBS) before plating for CFU count.

### Preparation of inoculum for infection of mice

*P. aeruginosa* PAO-1 was cultured in 5 ml TSB at 37 ℃ with shaking overnight. The culture was centrifuged 2,700 g for 15 minutes at 4 ℃ and the bacterial cell pellet was resuspended in 1 ml PBS. To create a persistent infection of the lung, the concentrated bacterial culture was mixed with 9 ml of liquid TSA pre-equilibrated at 50 ℃, and was mixed with heavy mineral oil to embed bacteria in agar beads to be used for the infection, using the procedure as described (Facchini et al. 2014). The challenge inoculum of *P. aeruginosa* was established by a pilot experiment to be 10^7^ CFU/ml for C57BL/6 mice.

### Persistent *P. aeruginosa* lung infection mouse model

C57BL/6 male mice (22-25 g, 6-8 week old) from Charles River were used for infection as described (Facchini et al. 2014). Before challenge, mice were anesthetized with ketamine (50 mg/ml) and xylazine (5 mg/ml) in 0.9 % NaCl administered at a volume of 0.002 ml/g body weight by intraperitoneal injection. After mice were fully anaesthetized, they were placed in supine position, followed by intra-tracheal instillation with 50 μl of agar beads containing bacterial suspension (10^7^ CFU/ml) to infect mice. A persistent lung infection that is difficult to heal is created due to the use of agar beads, mineral oil and stationary phase culture used for infection.

### Drug treatment in the mouse persistent lung infection

Based on a pilot experiment, at 3 day post-infection, mice would have the highest CFU count and establish a stable persistent lung infection model as shown in the previous study (Facchini et al. 2014) To evaluate different drugs and drug combinations on treating the persistent infection, the following groups of mice were used with each group having 5 mice per group: (1) drug free control (PBS); (2) cefuroxime (40 mg/kg) + gentamicin (30 mg/kg) treatment; (3) cefuroxime + clinafloxacin (40 mg/kg) treatment; (4) cefuroxime + gentamicin + levofloxacin (40 mg/kg) treatment; (5) clinafloxacin treatment; (6) cefuroxime + gentamicin + clinafloxacin treatment. Mice were treated daily intraperitoneally. After 7 days of treatment, the mice were sacrificed and the whole lung of each mouse was excised aseptically and homogenized in 1 ml PBS, and 100 μl of appropriately serial diluted lung homogenates were plated on TSA plate, followed by incubation at 37 ℃ overnight for CFU count.

## Results

### Use of SYBR Green I/PI assay to assess the viability of *P. aeruginosa*

The SYBR Green I/PI assay is a rapid and convenient method to assess bacterial viability under drug exposure at different time points (Feng et al. 2014). Thus, we first optimized the SYBR Green I/PI assay for use in determining *P. aeruginosa* viability as described (Feng et al. 2014). To generate a standard curve, we combined live and isopropyl-killed *P. aeruginosa* samples in ratios of 0:10, 1:4, 5:5, 4:1, 10:0 and after staining with SYBR Green I/PI, the green (live) and red (dead) fluorescence intensities of the samples were measured using a microplate reader (BioTek Instrument). A linear relationship (R^2^ values of 0.90561) between the ratios of green /red fluorescence and the percentages of live *P. aeruginosa* bacterial cells was established (Figure 1A). Fluorescence microscopy imaging confirmed the varying proportions of live and/or dead bacteria (Figure 1B).

**Figure 1.**
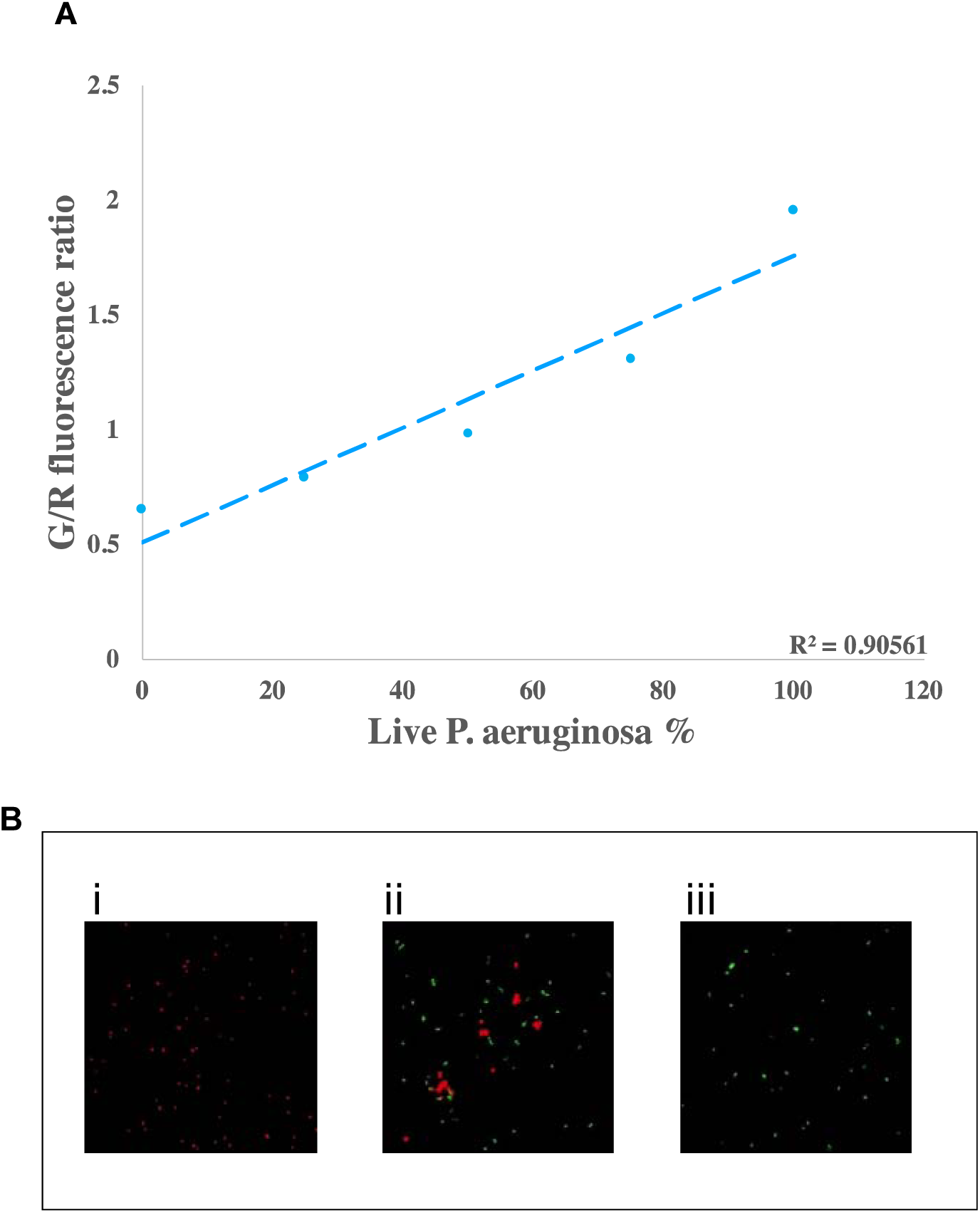
SYBR Green/PI staining can assess the viability of *P. aeruginosa* (A). A standard curve revealed linear relationship between the Green/Red fluorescence ratios from SYBR Green/PI viability assay and the percentage of live *P. aeruginosa* PAO-1 cells. The dead organisms were prepared with 70 % isopropyl alcohol and different proportions of live and dead cells were mixed and stained with SYBR Green I/PI and the green/red fluorescence ratios were measured by a microplate reader. **(B)** Representative microscope images of 0%, 50% and 100% of live *P. aeruginosa* cells using SYBR Green I/PI stain (200 X magnification). Green cells represent live cells and Red cells represent dead cells.

### Ranking of major classes of antibiotics for activity against stationary phase *P. aeruginosa*

To determine the relative activity of different classes of antibiotics against stationary phase *P. aeruginosa*, we performed drug exposure assays (all at 25 μM) with cell wall inhibitor (e.g. Cefuroxime), cell membrane inhibitor (Colistin), DNA synthesis inhibitor (e.g. Clinafloxacin), protein synthesis inhibitor (e.g. Gentamicin), RNA synthesis inhibitor (e.g. Rifampicin), Sulfa drug (e.g. Sulfamethoxazole), and Nitrofurantoin against *P. aeruginosa* stationary phase bacteria (Figure 2A). After 2 days of drug exposure, Clinafloxacin showed the highest activity in killing stationary phase bacteria, resulting in 0 CFU (Figure 2B). Cefuroxime and Colistin, cell wall and cell membrane inhibitors, respectively, and Sulfamethoxazole, had high activity against stationary phase bacteria compared with the drug free control. In contrast, Gentamicin, Rifampicin and Nitrofurantoin showed poor activity against the stationary phase *P. aeruginosa*.

**Figure 2.**
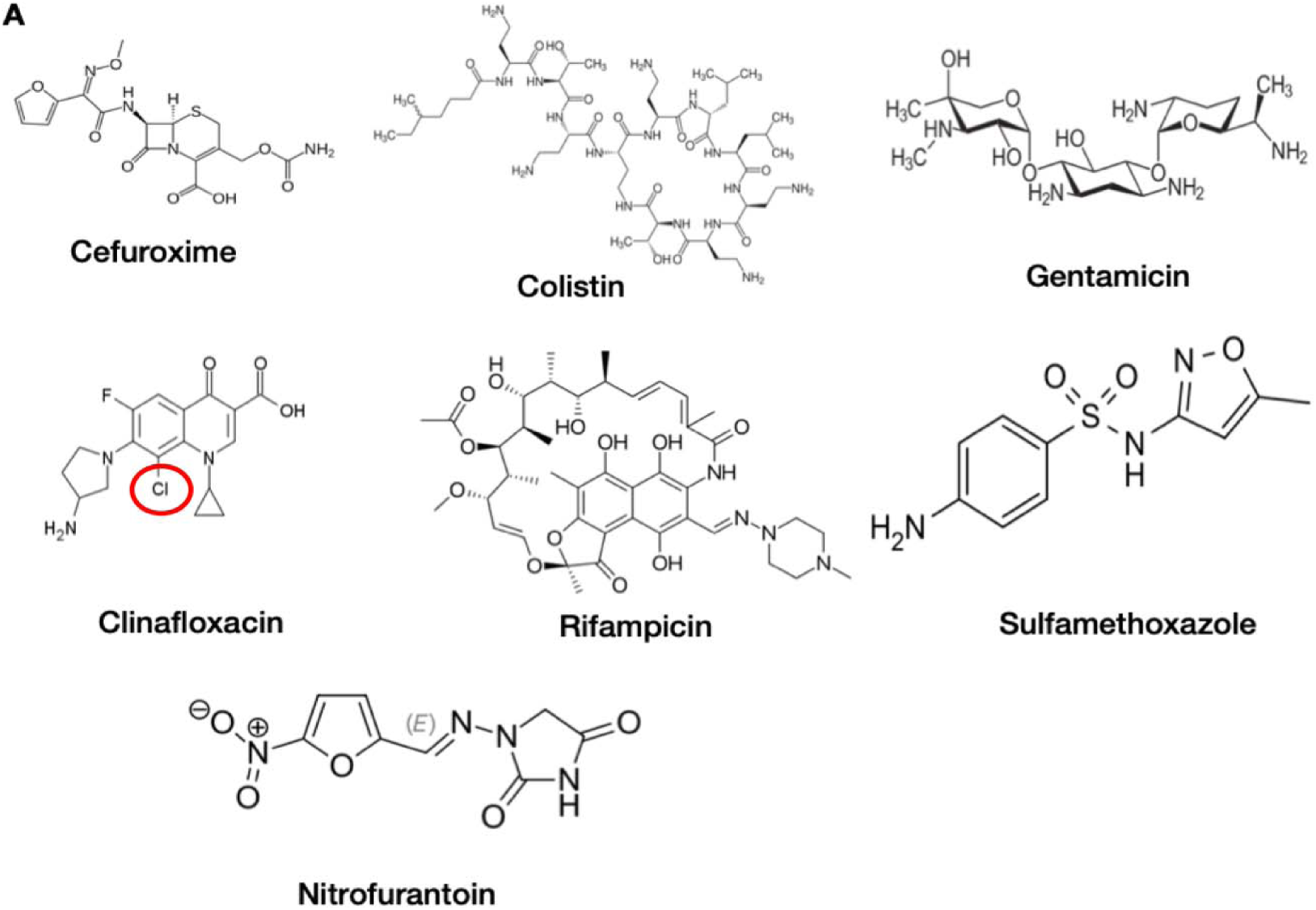
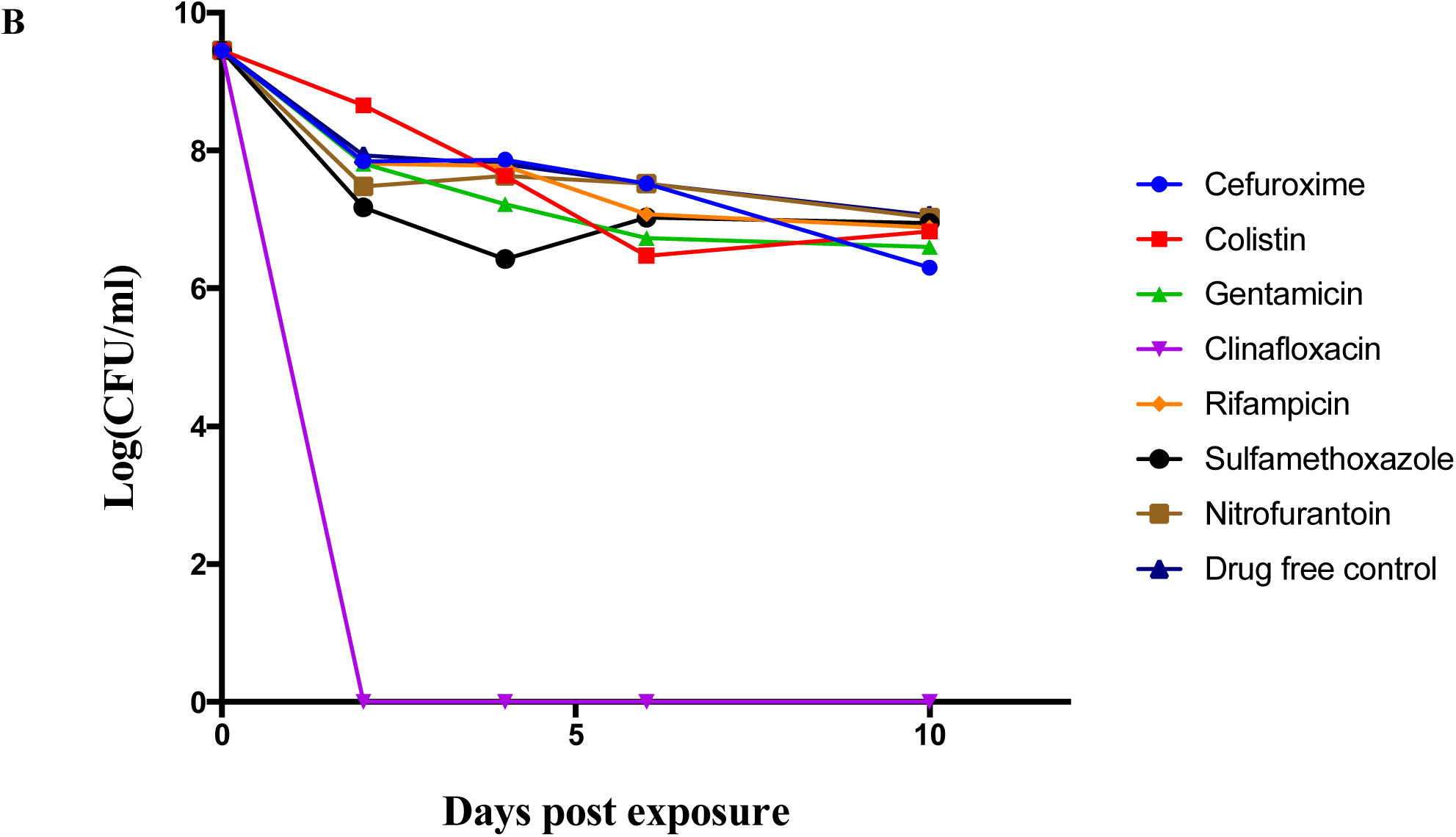
Ranking of the six classes of antibiotics by their activities against stationary phase *P. aeruginosa* determined by CFU assay. **(A)** Structures of the selected six different classes of antibiotics tested against stationary phase *P. aeruginosa* PAO-1 strain. **(B)** After 10 days of drug exposure (25 μM), the activity of each antibiotic against stationary phase *P. aeruginosa* was determined by CFU count.

Although the standard CFU assay can be used to evaluate the activity of antibiotics against stationary phase *P. aeruginosa*, we wanted to rank the same antibiotics by using the SYBR Green I/PI assay which is a more rapid method (Feng et al. 2014). Our results generated by SYBR Green I/PI viability assay correlated with findings from the CFU counting assay (Figure 3A). Cefuroxime and Colistin, Sulfamethoxazole and Clinafloxacin had among the highest activity against the stationary phase *P. aeruginosa.* The other three classes of antibiotics as represented by Gentamicin, Rifampicin and Nitrofurantoin had poor activity. We then calculated the residual viable bacterial cells after 10 day drug exposure through the regression equation (Table 1). After 10 days of treatment, Colistin killed the highest number of stationary phase bacteria, and Cefuroxime, Sulfamethoxazole and Clinafloxacin also showed remarkable effects with low percentages of residual viable bacterial cells remaining (Figure 3B). Like our results from CFU counting, the other three classes of antibiotics as represented by Gentamicin, Rifampicin and Nitrofurantoin killed fewer stationary phase bacteria with considerable numbers of residual viable cells remaining after treatment.

**Figure 3.**
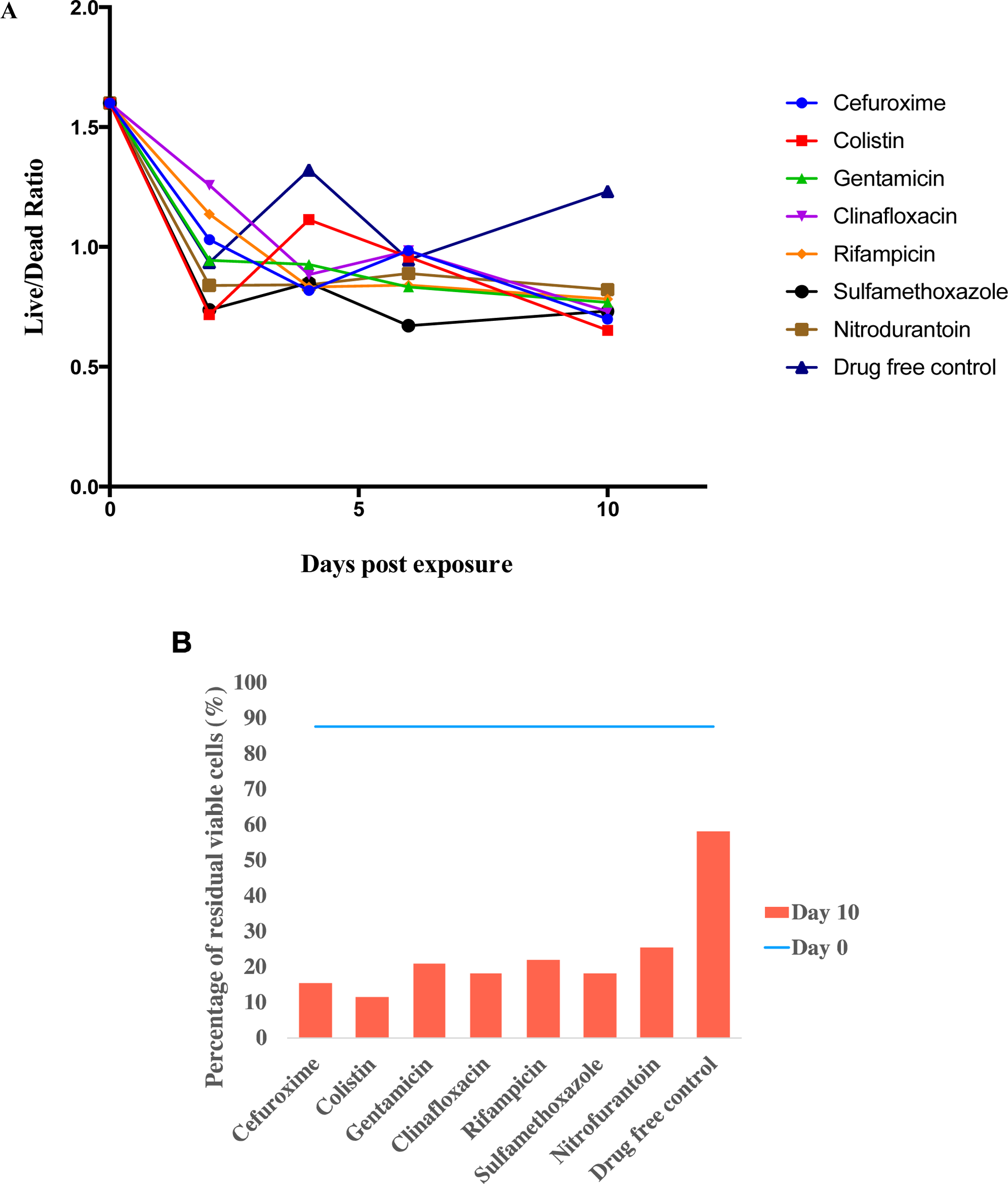
Ranking of the six classes of antibiotics by their activity against stationary phase *P. aeruginosa* determined by SYBR Green/PI viability assay. **(A)** After staining the bacterial culture with SYBR Green I/PI in the 96-well plate, the green/red fluorescence ratios were measured. SYBR Green/PI viability assay revealed drug activity results in concordance with the CFU assay. Cefuroxime and Colistin, Sulfamethoxazole and Clinafloxacin had good activity against stationary phase *P. aeruginosa.* **(B)** The percentage of residual viable cells after 10 day antibiotic treatment was calculated with the green/red ratios and the *P. aeruginosa* standard curve.

**Table 1.** Ranking of antibiotic activity from highest to lowest based on the amount of residual viable cells (Drug concentration: 25 μM)

| Drug Name | Residual viable cells (%) |
| --- | --- |
| Drug free control | 58.096 |
| Colistin | 11.696 |
| Cefuroxime | 15.456 |
| Clinafloxacin | 18.016 |
| Sulfamethoxazole | 18.096 |
| Gentamicin | 20.976 |
| Rifampicin | 22.096 |
| Nitrofurantoin | 25.296 |

\*The residual viable *P. aeruginosa* cells were calculated from a regression standard curve and equation from the SYBR Green I/PI assay as described by Feng et al. (2014).

Fluorescence microscopy analysis also confirmed that Cefuroxime, Colistin, Clinafloxacin and Sulfamethoxazole had the highest activities against stationary phase bacteria (Figure 4). At 10 day post-treatment, the majority of the bacterial cells were dead (as depicted in red PI stain) and the numbers of viable live cells (depicted as green in SYBR Green stain) were minimal. In contrast, stationary phase bacteria treated with Gentamicin, Rifampicin or Nitrofurantoin had more viable bacterial cells than dead cells, indicating their poor activity in killing stationary phase bacteria. Although these six classes of antibiotics all had activity against stationary phase bacteria compared to the drug free control, their relative activity against *P. aeruginosa* was quite different. From our CFU counts and our SYBR Green I/PI results, the six classes of antibiotics ranked from highest to lowest activity are as follows: clinafloxacin > colistin > gentamicin > cefuroxime > sulfamethoxazole > rifampicin > nitrofurantoin (Table 1).

**Figure 4.**
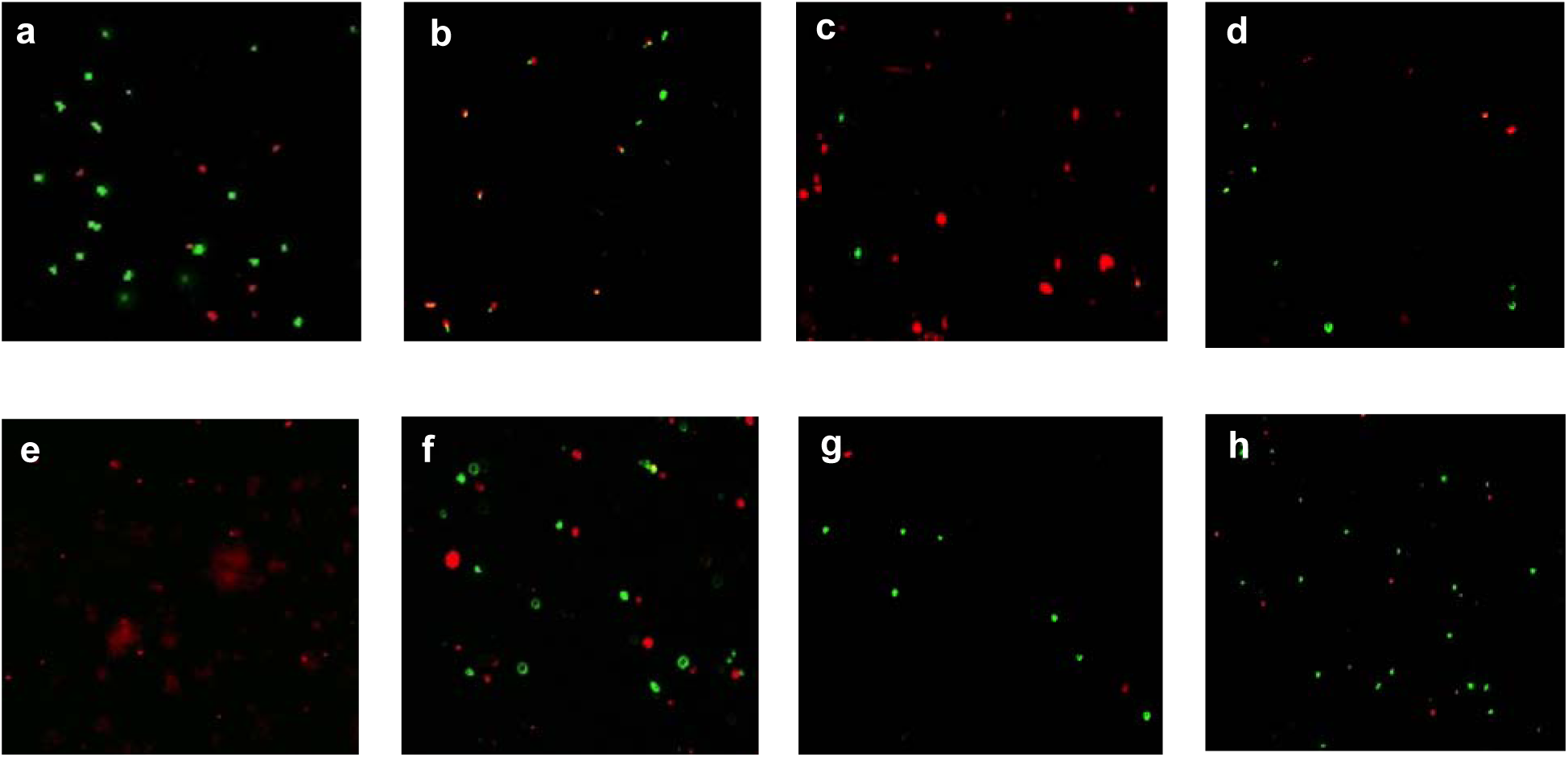
Fluorescence microscopy confirmation of stationary phase *P. aeruginosa* PAO-1 treated with six different classes of antibiotics (25 μM) followed by staining by SYBR Green I/PI assay (200X magnification). **(a)** Drug-free control, **(b)** Cefuroxime, **(c)** Colistin, **(d)** Gentamicin, **(e)** Clinafloxacin, **(f)** Rifampicin, **(g)** Sulfamethoxazole, and **(h)** Nitrofurantoin.

### Cefuroxime exhibits greater activity against stationary phase *P. aeruginosa* than other cell wall inhibitors

Cell wall inhibitors usually do not have good activity against persisters. Here, we found that cefuroxime at 25 μM reduced stationary phase bacteria from 10^9^ to 10^6^ CFU/ml (Figure 2B). To determine if it is the specific activity of cephalosporin cefuroxime against persisters, we tested other cell wall inhibitors such as amoxicillin and meropenem and found that compared to the dramatic persister killing activity of cefuroxime, amoxicillin and meropenem had limited activity against *P. aeruginosa* persisters as more than 10^7^ CFU/ml bacteria remaining after 7-day drug exposure (Figure 5A).

**Figure 5.**
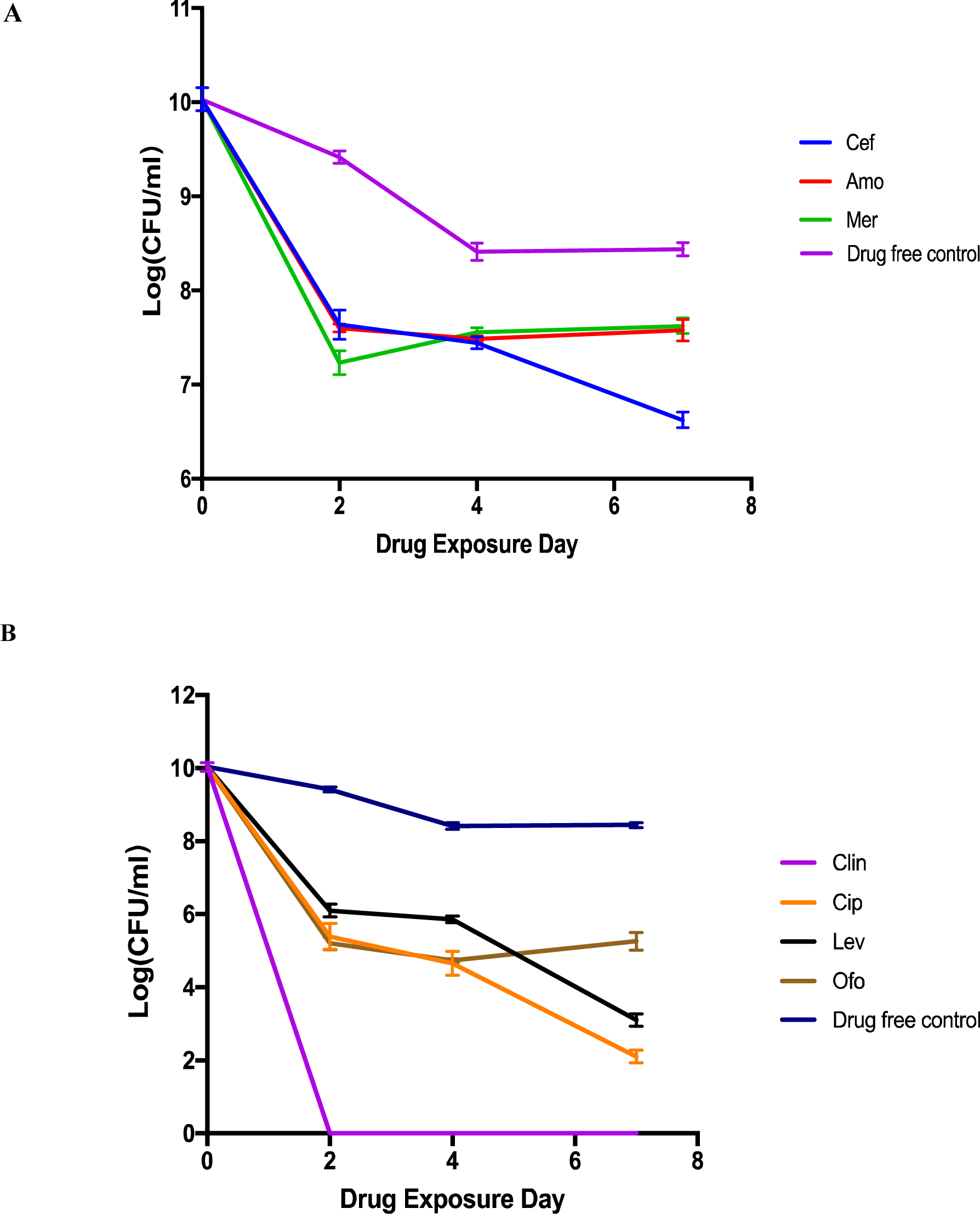
Comparison of relative activity of different cell wall inhibitors and different quinolone antibiotics against stationary phase *P. aeruginosa* by CFU assay. Stationary phase cultures of *P. aeruginosa* were exposed to different cell wall inhibitors (25 μM) (A) or different quinolone antibiotics (25 μM) for 0, 2, 4, 7 days when the CFU count was performed after washing and serial dilutions as described in Methods. Cefuroxime (Cef); Amoxicillin (Amo); Meropenem (Mer); Clinafloxacin (Clin); Ciprofloxacin (Cip); Levofloxacin (Lev); Ofloxacin (Ofo).

### Clinafloxacin exhibits the highest activity against stationary phase *P. aeruginosa* among different quinolone antibiotics

Since we found clinafloxacin had excellent activity against stationary phase bacteria (Figure 2B), we wanted to compare its activity with that of currently recommended quinolone antibiotics ciprofloxacin and levofloxacin in the treatment of *P. aeruginosa* infection(Johns Hopkins Hospital 2017), in terms of their activity against stationary phase *P. aeruginosa*. With the same 25 μM concentration, we found that only clinafloxacin could eradicate *P. aeruginosa* persisters only after 2 days by CFU count (Figure 5B). In contrast, ciprofloxacin and levofloxacin treatment still had 10^3^ and 10^4^ CFU/ml remaining, respectively.

### Relationship between the MIC values of antibiotics and their activity against stationary phase *P. aeruginosa*

Antibiotics with a low MIC value may have great activity against growing bacteria but may not have strong activity against stationary bacteria and vice versa in previous studies with *S. aureus* and *E coli* (Zhang 2005; Niu, Cui, Yee, et al. 2015; Niu, Cui, Shi, et al. 2015). Thus, we also tried to make a ranking among the six major classes of antibiotics (cell wall inhibitor, cell membrane disruptor, protein synthesis inhibitor, DNA synthesis inhibitor, RNA synthesis inhibitor, and anti-metabolite folate inhibitor) for their ability to kill growing log-phase bacteria. Based on the MIC values determined, our data showed that Colistin and Clinafloxacin had the lowest MIC values for *P. aeruginosa* PAO-1 strain (Table 2), which demonstrated that these two antibiotics can effectively kill both log phase and stationary phase bacteria. We also observed that Gentamicin was highly active against log phase bacteria but had low activity against stationary phase bacteria. Conversely, Cefuroxime and Sulfamethoxazole were less effective against growing PAO-1 bacteria, both with MICs above 16 μg/mL, but were effective in killing stationary phase bacteria (Tables 1 & 2). Rifampicin and Nitrofurantoin had low activity against both growing and stationary phase *P. aeruginosa*.

**Table 2.** Comparison of major classes of antibiotics and their respective activity against growing and stationary phase of *P. aeruginosa*.

| Class | Drug name | MIC<br>(µg/ml) | Cmax (µg/ml) | Activity against stationary phase<br>bacteria |
| --- | --- | --- | --- | --- |
| Cell wall<br>inhibitor | Cefuroxime | 16 | 4.1-7.0 | + |
| Cell membrane<br>disruptors | Colistin | 4 | 2.4 | +++ |
| Protein synthesis<br>inhibitor | Gentamicin | 1 | 5-12 | + |
| DNA synthesis<br>inhibitor | Clinafloxacin | 1 | 1.5-2.1 | +++ |
| RNA synthesis<br>inhibitor | Rifampicin | 16 | 8.2-11.7 | - |
| Sulfa drug | Sulfamethoxazole | 16 | 46.3 | ++ |
| Nitrofurantoin | Nitrofurantoin | >16 | 0.72 | - |

a Cmax values are derived from the literature.

### Development of drug combinations to eradicate *P. aeruginosa* biofilms *in vitro*

The bacterial cells within a biofilm structure are heterogeneous and contain persister cells that are tolerant to antibiotics (Williamson et al. 2012). Our previous work with *B. burgdorferi* biofilm-like structures indicated that a drug combination approach utilizing drugs active against growing bacteria and non-growing persister bacteria respectively is required to more effectively eradicate biofilm bacteria (Feng et al. 2016). Using the drugs that we have ranked with high activity against *P. aeruginosa* stationary phase persister cells, we tested the activity of various two-drug combinations (at clinically relevant Cmax concentrations) in killing stationary phase cells and biofilms. Our data demonstrate that only Clinafloxacin as a single drug could kill stationary phase cells completely (10^9^ CFU) after 4-day treatment, while single drugs such as colistin, cefuroxime, and gentamicin could kill only about 2-logs of stationary phase cells (Figure 6A). By adding Cefuroxime (a drug that has great activity killing growing bacteria) to the combination with anti-persister drug Clinafloxacin, the time to kill all stationary phase cells and biofilms was shortened from 4 days to 2 days. Compared to the clinical treatment cefuroxime + gentamicin + colistin, our designed drug combination cefuroxime + gentamicin + clinafloxacin could kill all biofilm (10^9^ CFU) after 4-day drug treatment, while single drugs or two drug combinations without clinafloxacin could kill only about 3-logs of biofilm cells and the two drug combinations with clinafloxacin could kill biofilm bacteria significantly to 10^2^ CFU/ml remaining (Figure 6B). In contrast, two-drug combinations such as colistin + gentamicin or cefuroxime + gentamicin, a combination currently used clinically, and even a three-drug combination cefuroxime + gentamicin + colistin only killed about 10^3^-10^4^ stationary phase and biofilm cells in 4 days, suggesting that a combination without an anti-persister drug (e.g. clinafloxacin) is not as effective. While the two-drug combination of colistin + gentamicin could also kill stationary phase cells by 10^8^ CFU after 4-day treatment, this combination was ineffective against biofilms.

**Figure 6.**
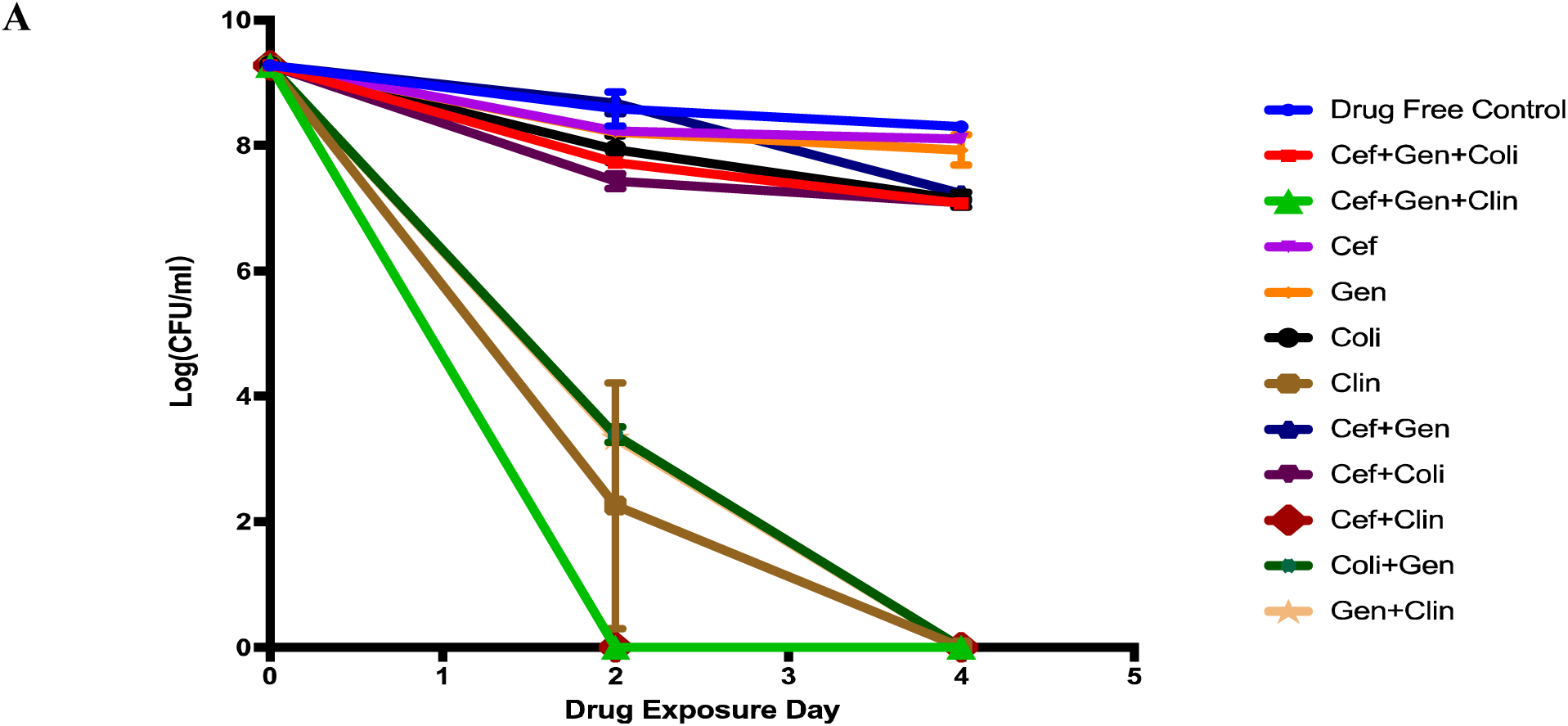
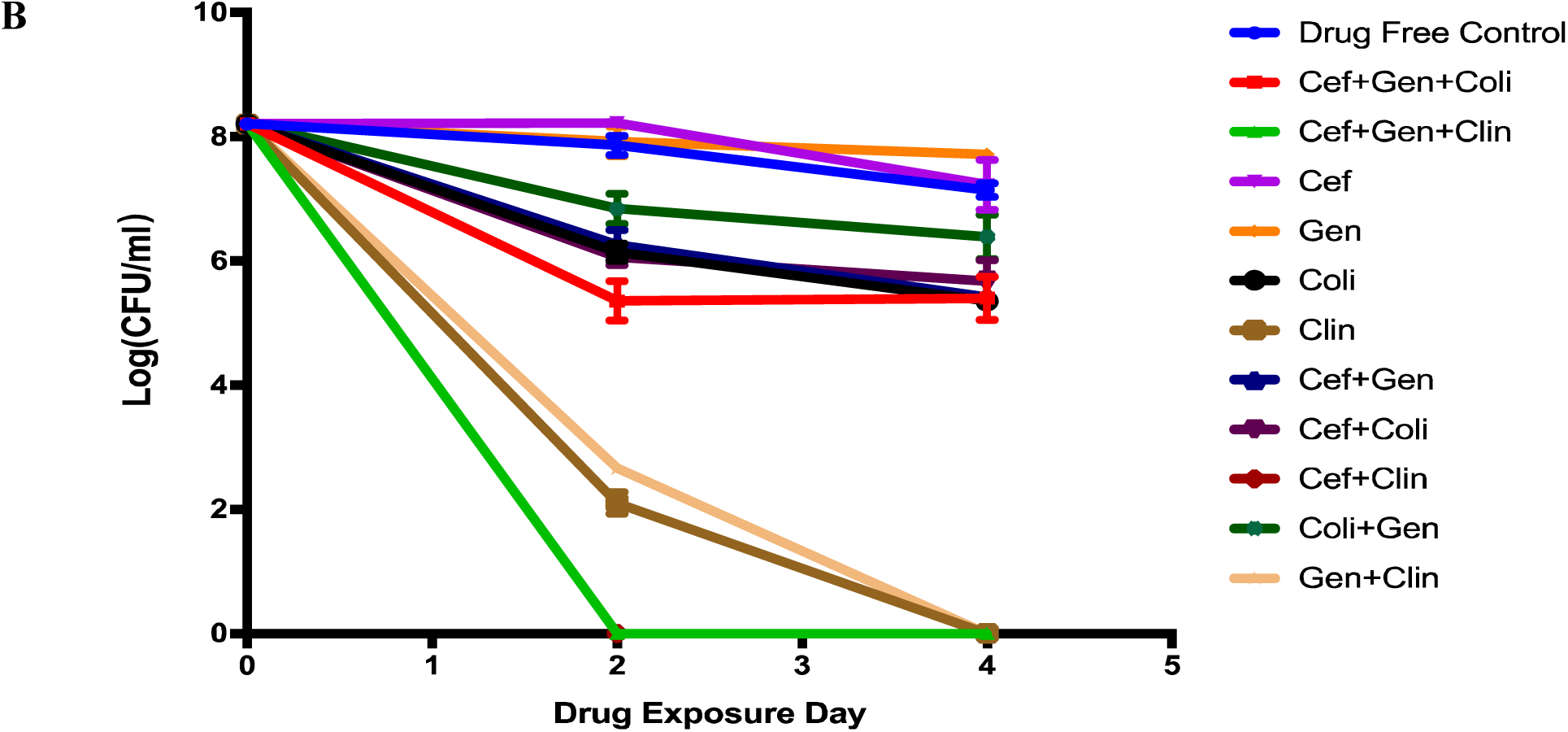
Comparison of the relative activity of different antibiotic combinations against stationary phase *P. aeruginosa* and biofilm bacteria by CFU assay. The effects of Clinafloxacin in drug combinations against stationary phase *P. aeruginosa* (A) and against *P. aeruginosa* biofilm model (B) were evaluated in drug exposure assays as described in Methods. CFU assay was performed on Day 0, 2, 4 after washing and serial dilutions. Cefuroxime (Cef): 5 μg/ml; Gentamicin (Gen): 10 μg/ml; Colistin (Coli): 5 μg/ml; Clinafloxacin (Clin): 1.5 μg/ml).

### Cefuroxime + Gentamicin + Clinafloxacin drug combination successfully eradicated persistent *P. aeruginosa* infection in cystic fibrosis model in mice

Since we found that cefuroxime + gentamicin + clinafloxacin completely eradicated stationary phase culture and biofilms in virto, we wanted to test if this triple persister drug regimen could also eradicate a persistent *P. aeruginosa* infection *in vivo*. To do so, we utilized a cystic fibrosis persistent *P. aeruginosa* infection model in mice as described (Facchini et al. 2014). Mice were infected with stationary phase *P. aeruginosa* PAO-1 mixed in beads through intra-tracheal instillation as described in Methods. After 3 day infection, bacteria reached peak at about 10^8^ CFU/g of the infected lungs when different treatments were started and continued for 7 days. There was a modest decrease of bacterial load (10^3^ fold) in the un-treated group after 7 days due to the host immune clearance of the bacterial infection(Facchini et al. 2014). After 7 day drug treatment, the current clinical recommended drug combination Cef + Gen only reduced the bacterial load from 10^8^ CFU/g to 10^4^ CFU/g (Figure 7). Clinafloxacin (Clin) alone, or Cef+Clin, reduced the bacterial load to 10^4^ or 10^3^ CFU/g, respectively, but were unable to completely clear the bacterial load in the lungs. In contrast, only Cef + Gen + Clin combination cleared all bacteria in the infected lungs. When we replaced Clin with levofloxacin (Lev) in the drug combination, we found that Cef + Gen + Lev combination was unable to clear the bacterial load and still had close to 10^4^ CFU/g remaining, indicating that the complete sterilization of the bacterial load is a property of Clinafloxacin in the context of combination with Cef and Gen. Thus, drug combination with clinafloxacin in this persistent lung infection model validated our *in vitro* drug combination results on biofilm bacteria.

**Figure 7.**
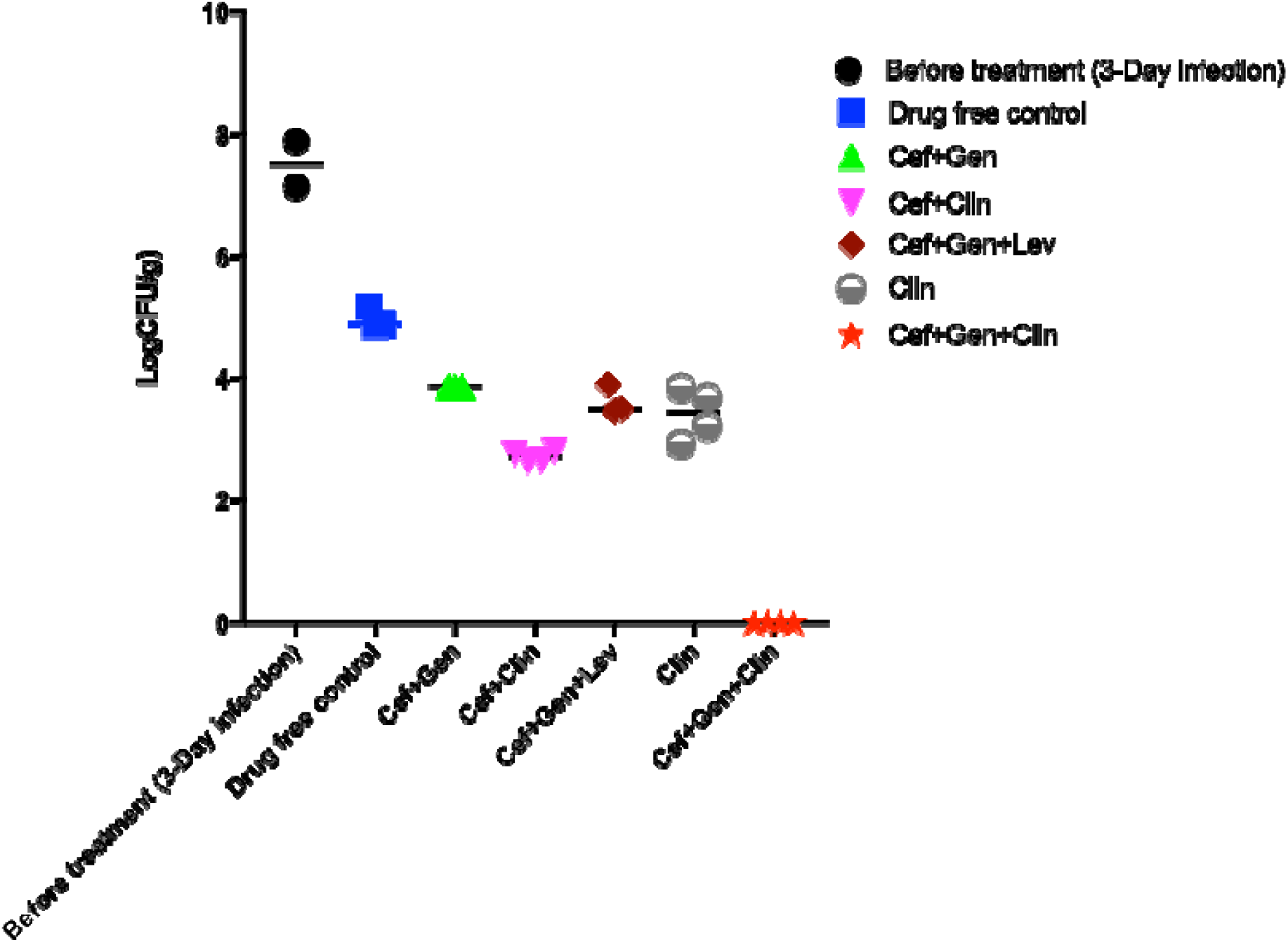
Cefuroxime + Gentamicin + Clinafloxacin drug combination successfully eradicated *P. aeruginosa* persistent infection in the mouse cystic fibrosis model. Persistent *P. aeruginosa* lung infection in mice was established as described in Methods. The treatment was started 3 days after infection and continued for 7 days when the CFUs from the lungs were determined. The treatment groups are as follows: before treatment group; drug-free control; Cef+Gen; Cef+Clin; Cef+Gen+Lev; Clin; Cef+Gen+Clin. Drug dosages used were Cef 40 mg/kg; Gen 30 mg/kg; Clin 40 mg/kg; Lev 40 mg/kg.

## Discussion

Clinically, it has been challenging to cure persistent infections like *P. aeruginosa* lung infections in cystic fibrosis patients. The difficulty to cure the persistent infections is thought to be mainly due to persister bacteria that are not effectively killed by the current antibiotics used to treat such infections. To address this hypothesis, here, we first ranked the six major classes of antibiotics for their activity against *P. aeruginosa* persisters in vitro and found a ranking order based on both CFU count and our SYBR Green/PI results: clinafloxacin > colistin > gentamicin > cefuroxime > sulfamethoxazole > rifampicin > nitrofurantoin (Figure 2B; Figure 3; Table 1). Among different cell wall inhibitors, cefuroxime showed greater activity to kill *P. aeruginosa* persisters than amoxicillin and meropenem (Figure 5A). Among different quinolone antibiotics, clinafloxacin killed all persister bacteria after 2 day drug exposure but other quinolones were not able to eradicate all persisters even after 7-day drug exposure (Figure 5B), suggesting that clinafloxacin could be a potential persister drug candidate for inclusion in drug combinations to kill persisters and biofilm bacteria. Based on the Yin-Yang model (Zhang 2014) utilizing drugs with activity against growing bacteria (MICs) and anti-persister activity results (Table 2), we designed a series of drug combinations to evaluate their ability to eradicate stationary phase or biofilm bacteria *in vitro*. Although the double drug combination of cefuroxime + clinafloxacin and gentamicin + clinafloxacin could eradicate persisters on day 4 treatment which has the same activity as the triple therapy cefuroxime + gentamicin + clinafloxacin (Cmax as concentration of each drug) and effectively killed stationary phase bacteria (Figure 6A), only the triple drug combination cefuroxime + gentamicin + clinafloxacin could kill all persisters after 2-day drug exposure in the *P. aeruginosa* biofilm model (Figure 6B). To further confirm the efficacy of the triple drug combination cefuroxime + gentamicin + clinafloxacin *in vivo*, we successfully established the *P. aeruginosa* lung persistent infection mouse model using stationary phase bacteria mixed with agar beads for intra-tracheal instillation infection. Using this persistent lung infection model, we were able to validate that clinafloxacin in combination with cefuroxime and gentamicin cleared all bacterial infection in the mouse lungs, whereas other single drugs or two drug combinations failed to eradicate the *P. aeruginosa* persistent lung infection (Figure 7).

Based on the Johns Hopkins Hospital Antibiotics Guideline (Johns Hopkins Hospital 2017), drugs that are included in routine treatment of *P. aeruginosa* infections range from β-lactam, aminoglycosides, fluoroquinolones to polymyxins. The majority of cystic fibrosis patients suffer from chronic infections of the airway with *P. aeruginosa* and the empiric treatment for *P. aeruginosa* currently uses two active agents from two different classes of antibiotics (Martinez et al. 2010). The two drug combination using high doses of synergistic antibiotic combinations (β-lactam + aminoglycoside) has been considered to improve outcomes of serious infections in immunocompromised hosts (Scribner et al. 1982). For multidrug resistant strains, colistin can be added to the above treatment (Florescu et al. 2012). However, in our study, we found that the current recommended treatment β-lactam + aminoglycoside (Cef + Gen) did not completely kill *P. aeruginosa* stationary phase and biofilm bacteria *in vitro* and failed to sterilize the lungs in the persistent lung infection in mice. In contrast, our persister drug Clinafloxacin in combination with Cef + Gen completely eradicated biofilm bacteria *in vitro* (Figure 6), and more importantly, the persistent lung infection in mice (Figure 7).

Our effective drug combination findings successfully apply the ‘Yin-Yang’ model to kill both replicating log phase bacteria (Yang) and also non-replicating persister bacteria(Yin)(Zhang 2014). In the Tuberculosis (TB) therapy, isoniazid (INH) is only active for growing mycobacteria, rifampin (RIF) is used to kill both growing and non-growing bacteria, and pyrazinamide (PZA) is added to kill exclusively non-growing persister bacteria in the treatment of TB (Zhang, Yew, and Barer 2012). The importance of PZA in killing persister TB bacteria and shortening the TB therpay without relapse is mainly recognized in the TB field (Zhang, Yew, and Barer 2012) and not appreciated in other persistent infections. We proposed to apply the PZA persister drug principle for the treatment of other infections (Zhang et al. 2013). This PZA principle has been shown to be valid in subsequent studies with different bacteria such as *B. burgdorferi* (Feng, Auwaerter, and Zhang 2015; Feng et al. 2016)*, E. coli* (Cui et al. 2016) and *S. aureus* (Yee R et al. to be published), both *in vitro* and *in vivo.* In this study, our findings with Clin + Cef +Gen combination both *in vitro* and *in vivo* further validate this PZA principle and the Yin-Yang model for developing more effective treatment of persistent infections, where Clin is an excellent drug targeting dormant persister bacteria (Yin) when used together with Cef and Gen which target growing bacterial populations for effective eradication of *P. aeruginosa* persistent lung infection (Figure 7). In our separate study with *S. aureus* pesistent skin infection model, we were also able to show that persister drug combinations utilizing the Yin-Yang model could achieve eradication of biofilm persistent infections (Yee, et al., to be published). Thus, we believe the PZA principle and Yin-Yang model can be applied for more effective treatment of other persistent infections in general.

There remain several areas for further investigation. First, the structural basis for the anti-persister activity in clinafloxacin fluoroquinolone drug is not clear but may be related to the unique chloride group in the quinolone ring (Figure 2). Further synthetic chemistry studies are needed to determine if the unique anti-persister activity is due to the chloride group and to further optimize the activity of clinafloxacin for anti-persister activity and to reduce its potential toxicity. Second, the mechanism of the unique anti-persister activity in clinfloxacin is not known but could be due to its activity on the bacterial cell membrane, as it is known that several persister drugs such as pyrazinamide (Zhang and Mitchison 2003), daptomycin (Feng, Auwaerter, and Zhang 2015), and colistin (Niu, Cui, Shi, et al. 2015; Cui et al. 2016), have activity on disrupting the bacterial membranes (Hurdle et al. 2011). Further studies are needed to understand how Clinafloxacin kills persisters more effectively than other fluoroquinolones. Third, despite the impressive anti-persister activity of clinafloxacin and its compassionate use for treatment of *Burkholderia cenocepacia* infection in a cystic fibrosis patient (Balwan et al. 2015), it is not an FDA approved drug because of its adverse drug reactions such as photosensitivity and hypoglycemia (Siami, LaFleur, and Siami 2002). It remains to be determined if the unique high anti-persister activity of clinafloxacin could outweigh its potential side effect in case of difficult to treat persistent infections. Thus, although our findings are encouraging, further studies are needed to address the above issues and its unique mechanism of anti-persister activity and its potential safety concerns in the future.

## Acknowledgement

YZ was supported in part by Cohen Foundation and Global Lyme Alliance.

